# *Acinetobacter baylyi* regulates type IV pilus synthesis by employing two extension motors and a motor protein inhibitor

**DOI:** 10.1101/2020.09.28.317149

**Authors:** Courtney K. Ellison, Triana N. Dalia, Catherine A. Klancher, Joshua W. Shaevitz, Zemer Gitai, Ankur B. Dalia

## Abstract

Bacteria use extracellular appendages called type IV pili (T4P) for diverse behaviors including DNA uptake, surface sensing, virulence, protein secretion, and twitching motility^1^. Dynamic extension and retraction of T4P is essential for their function, yet little is known about the mechanisms controlling these dynamics or the extent to which their regulation is conserved across bacterial species. Here, we develop *Acinetobacter baylyi* as a new model to study T4P by employing a recently developed pilus labeling method^2,3^. Our findings overturn the current dogma that T4P extension occurs through the action of a single, highly conserved motor, PilB, by showing that T4P synthesis in *A. baylyi* is dependent on an additional, phylogenetically distinct motor, TfpB. Furthermore, we uncover an inhibitor of T4P extension that specifically binds to and inhibits PilB but not TfpB. These results expand our understanding of T4P regulation and highlight how inhibitors might be exploited to inhibit T4P synthesis.

## Introduction

T4P are thin, proteinaceous appendages that are broadly distributed throughout bacteria and archaea^4^. T4P are composed primarily of major pilin protein subunits that are polymerized or depolymerized through the activity of ATPases to mediate fiber extension and retraction, respectively^5^. T4P are subdivided into subcategories (generally T4aP, T4bP, and T4cP) based on protein homology and pilus function, with T4aP being the best-characterized. In the T4aP systems found in Gram-negative organisms, polymerization and depolymerization of pilins occurs through interactions of the extension ATPase PilB or retraction ATPase PilT with the integral inner membrane platform protein PilC (Supplemental figure 1a). The growing fiber spans an alignment complex from the inner membrane through the periplasm composed of PilNOP to exit through the PilQ outer membrane secretin pore. Dynamic cycles of T4P extension and retraction are critical for the diverse processes that these structures mediate including twitching motility^6^, surface sensing^2,7^, virulence^8,9^, and DNA uptake^10,11^.

*Acinetobacter* species like *A. baumannii* and *A. nosocomialis* have emerged to become an urgent medical threat due to their prevalence in hospital-acquired infections and their capacity to acquire antibiotic resistance genes; a process that is achieved in part by natural transformation through T4aP-mediated DNA uptake^12,13^. *Acinetobacter baylyi* is the most naturally transformable species reported to date^14^, with up to 50% of cells undergoing natural transformation in laboratory conditions, making it an ideal candidate to study T4P-mediated DNA uptake and natural transformation.

## Results

To study *A. baylyi* T4P, we applied a recently-developed labeling method^2,3^ by targeting the major pilin, ComP, for cysteine substitution and subsequent labeling with thiol-reactive maleimide dyes. Maleimide-labeling of the functional *comP*^T129C^ strain (Supplemental figure 1b, 1c) revealed external T4P filaments as seen in other species using this method^3^. T4P in *A. baylyi* appear much shorter than those found in other species like *V. cholerae*, and they localize close together in a line along the long axis of the cell (Supplemental figure 1c). Incubation of cells with fluorescently-labeled DNA resulted in co-localization of DNA with T4P, which is consistent with the essential role of T4P-DNA binding during natural transformation in *Vibrio cholerae*^11^ (Figure 1a). We reasoned that natural transformation could be used to screen for other factors that regulate T4P synthesis in *A. baylyi*. To that end, we performed a high throughput transposon-sequencing screen (Tn-seq)^15^ to identify genes required for natural transformation. In this screen, loss of known T4P-related genes resulted in negative selection, indicating that they were critical for natural transformation as expected (Figure 1b). To validate our Tn-seq results, we made in-frame deletions of representative genes from each T4P-encoding operon, as well as genes known to be essential for natural transformation that act downstream of T4P. Mutations in the T4P platform protein gene *pilC*, the pilus regulatory gene *pilY1*, the outer membrane secretin gene *pilQ* and the retraction ATPase gene *pilT* all showed a marked reduction in natural transformation, with most mutants exhibiting transformation rates below our limit of detection (Figure 1c). T4P-labeling of *pilC, pilY1*, and *pilQ* mutants revealed no visible T4P fibers (Figure 1d). Mutations in genes that act downstream of T4P, including the periplasmic DNA-binding protein gene *comEA*, the inner membrane DNA-transporter protein gene *comEC*, and the DNA-recombination helicase gene *comM* likewise resulted in a reduction in transformation frequency, however, these strains still produced T4P fibers as expected (Figure 1c, d).

**Figure 1.**
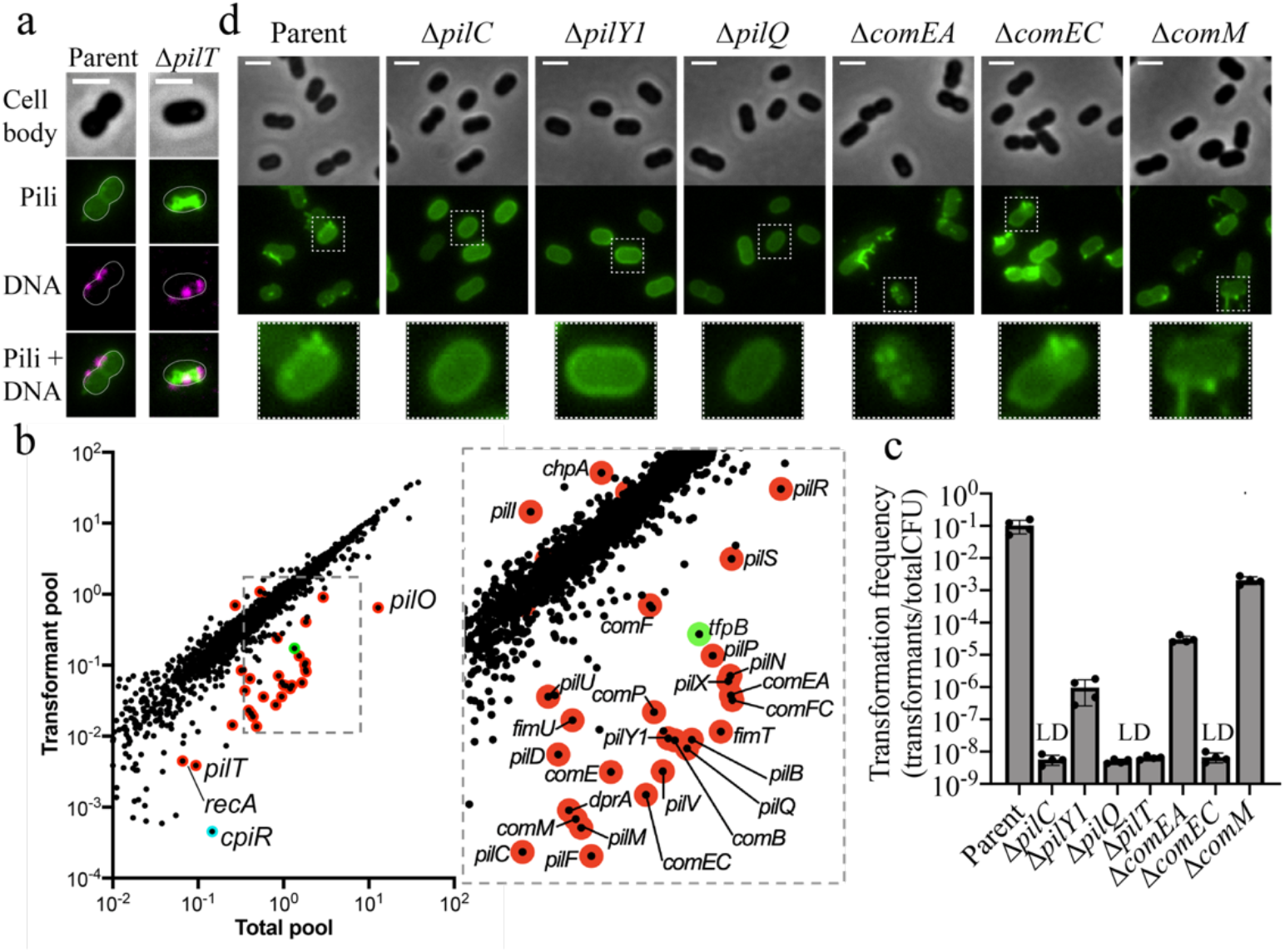
Tn-seq reveals factors important for natural transformation and T4P synthesis. (**a**) Representative images of cells with AF488-mal labeled pili incubated with fluorescently-labeled DNA. (**b**) Visual representation of Tn-seq screen showing the relative abundance of each gene in the “total pool” of transposon mutants compared to the “transformant pool” recovered after natural transformation. Known T4P structural/regulatory genes are outlined in red. *tfpB* is outlined in green, and *cpiR* is outlined in cyan. (**c**) Natural transformation assays of indicated strains. Each data point represents a biological replicate and bar graphs indicate the mean ± SD. LD, limit of detection. (**d**) Representative images of strains from natural transformation assays shown in **c** labeled with AF488-mal. Zoomed-in images of representative single cells from each strain are outlined in dashed boxes and shown below. Scale bars, 2 μm.

We next sought to determine whether any of the previously uncharacterized genes from our Tn-seq screen reduced natural transformation in *A. baylyi* by affecting T4P synthesis. Tn-seq revealed that mutations in *pilB*, the canonical extension ATPase gene that is typically cotranscribed with *pilC* and the pre-pilin peptidase, *pilD*, resulted in reduced natural transformation (Figure 1b). We also found an additional *pilB* homologue that likewise exhibited lower rates of transformation (Figure 1b). We named this new PilB homologue TfpB for “<u>t</u>ype <u>f</u>our <u>p</u>ilus PilB-like protein” because although it has homology to PilB, it does not exhibit the same gene synteny as *pilB* genes that are typically co-transcribed with their cognate inner membrane platform gene *pilC*. Surprisingly, deletion of the canonical *pilB* did not ablate transformation as would be expected based on homology to other T4P^10^ (Figure 2a). Mutation of *tfpB* reduced transformation rates to a level that was similar to the *pilB* mutant; however, natural transformation was undetectable in the *pilB tfpB* double mutant (Figure 2a).

**Figure 2.**
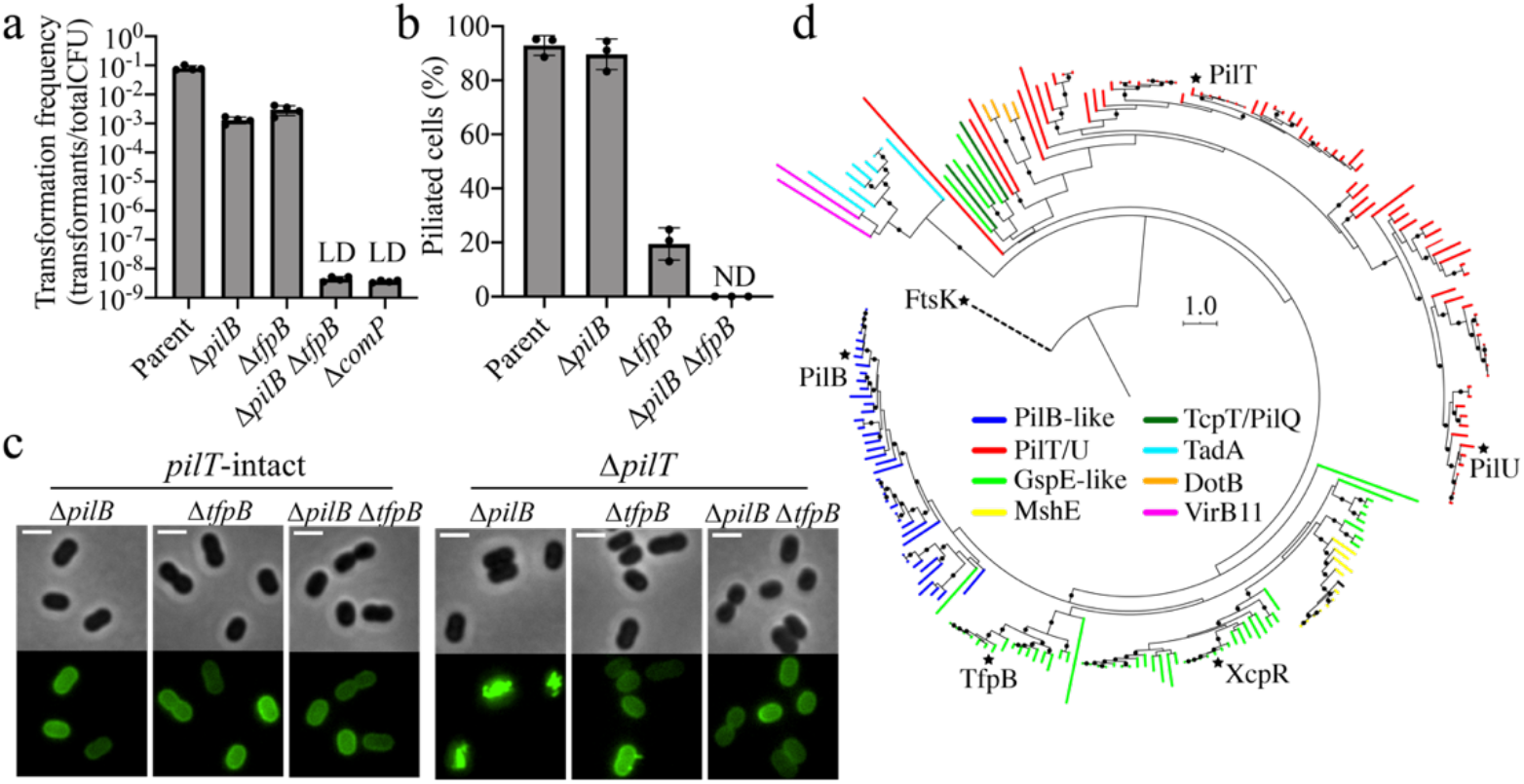
TfpB is a phylogenetically distinct PilB homologue that is required for efficient T4P extension in *A. baylyi*. (**a**) Natural transformation assays of indicated strains. Each data point represents a biological replicate and bar graphs indicate the mean ± SD. LD, limit of detection. (**b**) Percent of piliated cells in *pilT* mutant populations of indicated strains. Each data point represents an independent, biological replicate and bar graphs indicate the mean ± SD. For each biological replicate, a minimum of 70 total cells were assessed. ND, no pili detected. (**c**) Representative images of indicated strains labeled with AF488-mal with background fluorescence subtracted. Scale bars, 2 μm. (**d**) A rooted phylogeny of TfpB homologues found among Gammaproteobacteria. Branches are colored according to protein annotations in IMG, and nodes with bootstrap values greater than or equal to 70% are indicated by black circles. Black stars are at the tips of branches representing *A. baylyi* proteins with indicated protein names.

Deletion of *pilT* prevents T4P retraction^16^ and may thus reveal more subtle effects on pilus extension. T4P labeling in *pilB*, *tfpB*, and *pilB tfpB* mutants revealed that all three mutants were defective in T4P synthesis while *pilT* was intact (Figure 2c). In a *pilT* deletion background, *pilB* mutant populations had similar numbers of cells producing T4P compared to the Δ*pilT* parent while, surprisingly, *tfpB* mutants were highly defective in T4P synthesis (Figure 2b, c). The *pilB tfpB pilT* triple mutant produced no detectable T4P fibers (Figure 2b, c). Together, these results indicate that both PilB and TfpB are essential for efficient T4P extension in *A. baylyi*, with TfpB playing the dominant role.

Phylogenetic analysis of extension ATPase homologues found among Gammaproteobacteria^4,17^, which include members of the PilT/VirB11 family of secretion ATPases, revealed that TfpB clusters with a group of proteins that are phylogenetically distinct from the canonical PilB ATPase (Figure 2d, Supplemental figure 2). PilB and TfpB proteins are as divergent from one another as PilB and the type II secretion system proteins XcpR/GspE, or PilB and the MshE motors that drive mannose-sensitive haemagglutinin (MSHA) pilus synthesis, suggesting that TfpB evolved as a functionally divergent class of proteins that is distinct from canonical PilB extension motors. TfpB is highly conserved in other *Acinetobacter* species (Figure 2d, Supplemental figure 2), implying that use of multiple extension motors may be prevalent in the *Acinetobacter* clade. These results suggest that proteins that cluster with TfpB may play similar functions in other bacterial species and that multiple extension ATPases may be a common feature of diverse T4P.

In addition to revealing the importance of TfpB in T4P synthesis, our Tn-seq results also uncovered an uncharacterized transcriptional regulator belonging to the XRE-family of transcriptional repressors that is critical for natural transformation (Figure 1b, Figure 3a). Deletion of this regulator, named *cpiR* for <u>c</u>ompetence <u>p</u>ilus <u>i</u>nhibition <u>r</u>epressor, resulted in a 100-fold reduction in transformation (Figure 3b). We hypothesized that CpiR repressed a factor that inhibits natural transformation. Transcriptional repressors are often transcribed immediately adjacent to the genes they repress. We thus deleted the upstream gene, named *cpiA* for competence pilus inhibition actuator (Figure 3a), and found that transformation frequency was restored in the *cpiR cpiA* double mutant background (Figure 3b). Expression of *cpiA* under the control of an IPTG-inducible promoter (P*tac*) was sufficient to reduce transformation frequency even in the presence of *cpiR*, further suggesting that CpiR represses *cpiA* transcription. Furthermore, we found that CpiR was sufficient to repress the *cpiA* promoter (using a P_*cpiA*_-GFP reporter) in the heterologous host *Vibrio cholerae*, which suggests that CpiR is a direct repressor of P_*cpiA*_ (Figure 3c). Finally, we found that CpiA protein is only produced in a *cpiR* mutant (Figure 3d). These results demonstrate that under lab conditions, CpiR represses *cpiA* transcription to allow for high rates of natural transformation; however, in the absence of CpiR, CpiA is produced and natural transformation is inhibited ~100-fold.

**Figure 3.**
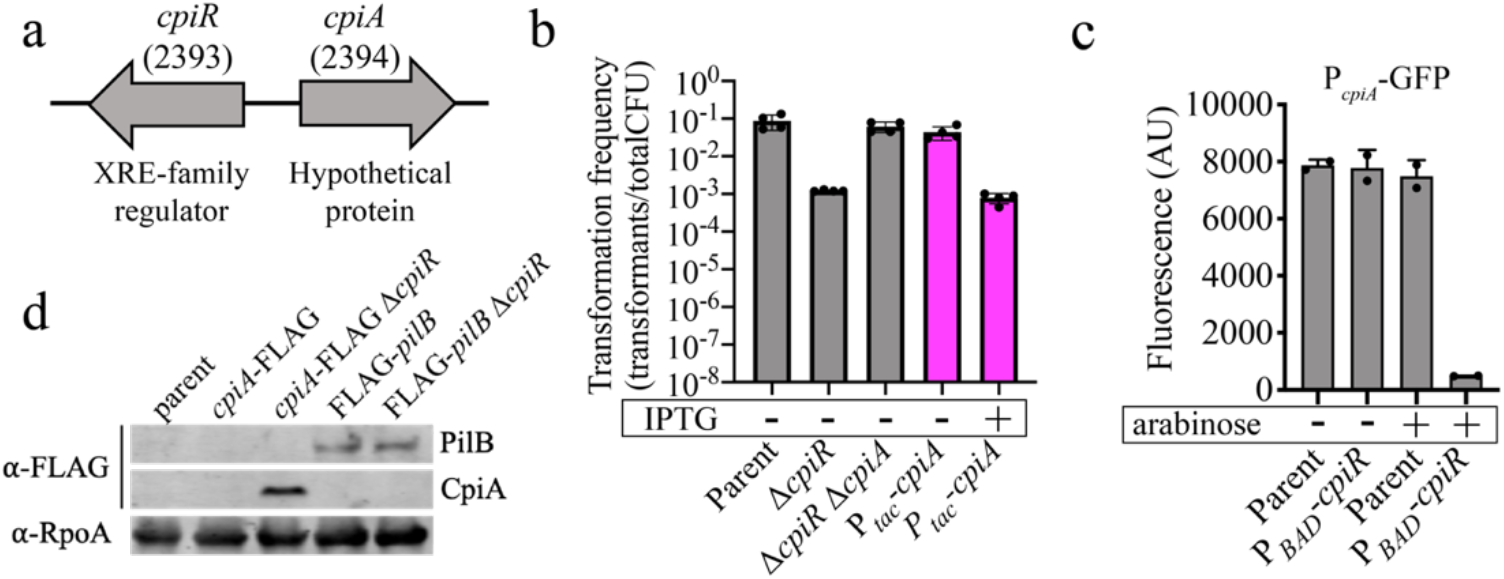
Natural transformation is regulated by the inhibitor protein, CpiA, whose expression is controlled by the transcriptional repressor, CpiR. (**a**) Schematic of *cpiRA* locus organization. Numbers in parentheses are ACIAD numbers associated with each gene. (**b**) Natural transformation assays of the indicated strains performed with or without 100 μM IPTG added as indicated. Each data point represents a biological replicate and bar graphs indicate the mean ± SD. Pink bars denote a strain where *cpiA* is expressed at an ectopic location under the control of an IPTG-inducible promoter. (**c**) Fluorescence intensity of *V. cholerae* strains harboring a P_*cpiA*_-GFP reporter and a P_*BAD*_-cpiR ectopic expression construct as indicated when grown with or without 0.2% arabinose as indicated. Each data point represents a biological replicate and bar graphs indicate the mean ± SD. (**d**) Western blot showing CpiA-FLAG production in the indicated strains and PilB levels when CpiA is expressed. RpoA was detected as a loading control.

CpiA lacks primary sequence or structural homology to any proteins or domains of known function^18,19^, so we next sought to determine the mechanism of transformation inhibition by CpiA. Labeling of T4P in a *cpiR* mutant (i.e. when CpiA is expressed) revealed that cells were deficient in T4P synthesis (Figure 4a). This defect was dependent on *cpiA* because piliation was restored in the *cpiR cpiA* double mutant (Figure 4a). Deletion of *pilT* restored T4P synthesis in the *cpiR* mutant (i.e. the *cpiR pilT* double mutant) (Figure 4a, b). This phenotype was reminiscent of the restoration of piliation observed in the *pilB pilT* double mutant, so we hypothesized that CpiA may regulate natural transformation by inhibiting PilB.

**Figure 4.**
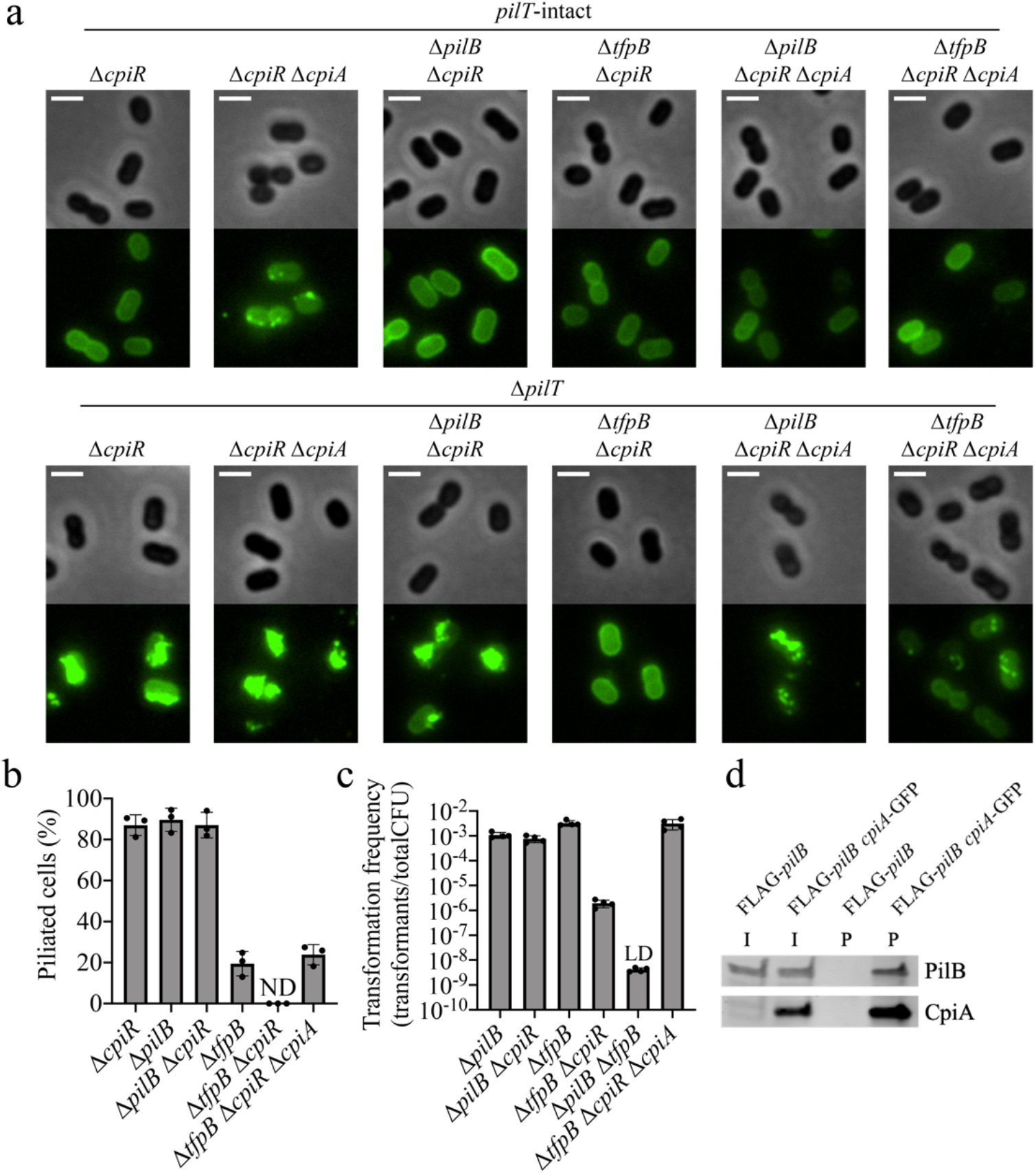
CpiA inhibits PilB activity. (**a**) Representative images of indicated strains labeled with AF488-mal with background fluorescence subtracted. Scale bars, 2 μm. (**b**) Percent of piliated cells in *pilT* mutant populations of indicated strains. Each data point represents an independent, biological replicate and bar graphs indicate the mean ± SD. For each biological replicate, a minimum of 70 total cells were assessed. ND, no pili detected. (**c**) Natural transformation assays of indicated strains. Each data point represents a biological replicate and bar graphs indicate the mean ± SD. LD, limit of detection. (**d**) Western blot showing coimmunoprecipitation experiments where CpiA-GFP was used as the bait protein to test for interaction with FLAG-PilB as the prey protein in Δ*cpiR* mutant backgrounds. I, input sample; P, pulldown (coimmunoprecipitation) sample.

To test if CpiA functions by inhibiting PilB, we made *pilB cpiR* and *tfpB cpiR* double mutants. T4P synthesis was unaffected in *pilB cpiR pilT* (where TfpB is the sole extension ATPase present), while no T4P were detected in *tfpB cpiR pilT* (where PilB is the sole extension ATPase present) (Figure 4a, b). The defect in T4P synthesis in the *tfpB cpiR pilT* strain was restored when *cpiA* was deleted in this background, demonstrating that CpiA acts as a PilB inhibitor to control natural transformation (Figure 4a, b). Transformation frequency assays corroborated these results by demonstrating that deletion of *cpiR* only inhibited transformation in the *tfpB* mutant background (where PilB is the sole extension ATPase) and not the *pilB* mutant background (where TfpB is the sole extension ATPase) (Figure 4c). Immunoblotting revealed that CpiA does not affect PilB production or stability (Figure 3d), and we thus hypothesized that CpiA may interact with PilB to disrupt its function. Coimmunoprecipitation experiments with functional fusion proteins (Supplemental figure 3) revealed that CpiA and PilB interact with each other, and that their interaction is not dependent on the pilus machinery proteins PilMNOPQ (Figure 4d and Supplemental figure 4), ruling out indirect binding of CpiA to PilB through other machinery components required for T4P synthesis. CpiA and TfpB do not interact (Supplemental figure 4), consistent with our genetic evidence that CpiA specifically inhibits PilB and not TfpB (Figure 4a-c). Together, these data suggest that CpiA directly binds to and inhibits PilB.

## Discussion

Environmental conditions play an integral role in regulating how bacteria respond to and interact with their surroundings. The mechanisms of T4P synthesis regulation identified here likely reflect how environmental conditions influence bacterial physiology. The acquisition of the additional extension motor TfpB and the PilB-specific inhibitory protein CpiA likely resulted from a need to modulate T4P extension under different environmental conditions. Bacteria generally limit T4P synthesis to environments where they provide a selective advantage. For example, in *V. cholerae*, the production of competence T4aP requires chitin and quorum sensing^20^, which are environmental conditions that cells experience when they are likely to encounter other bacterial cells for horizontal gene transfer; while the toxin-coregulated T4bP (TCP) required for intestinal colonization are produced in response to host-specific cues^21^. In *A. baylyi*, both PilB and TfpB are required for T4P extension and, consequently, efficient natural transformation. While CpiA is not expressed under laboratory conditions (due to CpiR repression), it is possible that different environmental conditions derepress *cpiA* to decrease pilus activity.

Secretion ATPases like PilB belong to a broadly distributed class of proteins that play essential roles in diverse microbial behaviors, yet there are few known inhibitors of these proteins. The ones that are characterized are encoded by phages that use T4P to infect cells, and it is speculated that PilB inhibition by these proteins prevents superinfection by other T4P-dependent phages^22,23^. In *Acinetobacter baylyi, cpiA* is encoded on the chromosome. Thus, it is tempting to speculate that CpiA was acquired after phage infection of an ancestral strain and co-opted for regulation of T4P activity. A better understanding of secretion ATPase function and the evolution of mechanisms by which they can be inhibited may enable the development of tools to control their activity. Precise control of T4P synthesis may provide a means to manipulate behaviors that require T4P like biofilm formation, virulence, and natural transformation, which are clinically relevant in diverse pathogens.

## Methods

### Bacterial strains and culture conditions

*Acinetobacter baylyi* strain ADP1 was used throughout this study. For a list of strains used throughout, see Supplemental Table 1. *A. baylyi* cultures were grown at 30 °C in Miller lysogeny broth (LB) medium and on agar supplemented with kanamycin (50 μg/mL), spectinomycin (60 μg/mL), gentamycin (30 μg/mL), and/or chloramphenicol (30 μg/mL), rifampicin (30 μg/mL), zeocin (100 μg/mL), and/or streptomycin (10 μg/mL) as appropriate.

### Construction of mutant strains

Mutants in *A. baylyi* were made using natural transformation as described previously^16^. Briefly, mutant constructs were made by splicing-by-overlap (SOE) PCR to stich 1) ~3 kb of the homologous region upstream of the gene of interest, 2) the mutation where appropriate (for deletion by allelic replacement with an antibiotic resistance cassette, or the fusion protein), and 3) the downstream region of homology. For a list of primers used to generate mutants in this study, see Supplemental Table 2. The upstream region was amplified using F1 + R1 primers, and the downstream region was amplified using F2 + R2 primers. All antibiotic resistance (AbR) cassettes were amplified with ABD123 (ATTCCGGGGATCCGTCGAC) and ABD124 (TGTAGGCTGGAGCTGCTTC). Fusion proteins were amplified using the primers indicated in Supplemental Table 2. In-frame deletions were constructed using F1 + R1 primer pairs to amplify the upstream region and F2 + R2 primer pairs to amplify the downstream region with ~20 bp homology to the remaining region of the downstream region built into the R1 primer and ~20 bp homology to the upstream region built into the F2 primer. SOE PCR reactions were performed using a mixture of the upstream and downstream regions, and middle region where appropriate using F1 + R2 primers. SOE PCR products were added with 50 μl of overnight-grown culture to 450 μl of LB in 2 ml round-bottom microcentrifuge tubes (USA Scientific) and grown at 30 °C rotating on a roller drum for 1-3 hours. For Ab^R^-constructs, transformants were serially diluted and plated on LB and LB + antibiotic. For in-frame deletions and protein fusion constructs, after the three-hour incubation, cells were incubated with 10 μl of DNaseI (New England Biolabs) for 5 min at room temperature before 450 μl of LB was added. Transformations were grown for an additional hour before cells were diluted and 100 μl of 10^-6^ dilution was plated on LB plates. In-frame deletions were confirmed by PCR using primers ~150 bp up and downstream of the introduced mutation, and fusions were confirmed by sequencing.

Mutants in *V. cholerae* were made by chitin-induced natural transformation exactly as previously described^24,25^. SOE products were generated exactly as described above. The chromosomally integrated P_*BAD*_ ectopic expression construct is described in detail in ref ^26^.

### Natural transformation assays

Assays were performed exactly as previously described^27^. Briefly, strains were grown overnight in LB broth at 30°C rolling. Then, ~10^8^ cells were subcultured into fresh LB medium and 100 ng of tDNA was added. In this study, a ΔACIAD1551::Spec^R^ PCR product (with 3kb arms of homology) was used as the tDNA. Reactions were incubated with end-over-end rotation at 30°C for 5 hours and then plated for quantitative culture on spectinomycin plates (to quantify transformants) and on plain LB plates (to quantify total viable counts). Data are reported as the transformation frequency, which is defined as the (CFU/mL of transformants) / (CFU/mL of total viable counts).

### Tn-seq Analysis

Transposon mutant libraries of *A. baylyi* were generated by electroporation of cells with pDL1093, a vector that allows for Kan^R^ mini-Tn10 mutagenesis^28,29^, and plating on kanamycin plates. Approximately 100,000 Tn mutants were scraped off of plates and pooled to generate the input transposon mutant library. This mutant library was then subjected to natural transformation in three separate replicates exactly as described above using three different sources of tDNA: a ΔACIAD1551::Spec^R^ PCR product, an RpsL K43R (Sm^R^) PCR product, or an RpoB PCR product from a spontaneous rifampicin resistant mutant. Following incubation with tDNA, reactions were split and outgrown in selective medium overnight (i.e. with the appropriate antibiotic added to select for transformants = “transformant pool”) or grown overnight in nonselective medium (i.e. in plain LB medium = “total pool”). Sequencing libraries of the transposon-genomic junctions were generated for the Illumina platform using HTM-PCR exactly as previously described^29,30^. Sequencing data were mapped to the *V. cholerae* N16961 genome^31^ and the relative abundance of Tn insertions in all samples was determined on the Galaxy platform^32^. Comparative analyses for Tn-seq were performed treating the “transformant pool” as the output and the “total pool” as the input to determine genes that were over- and under-represented following selection for transformants. Gene fitness was only assessed if a gene contained at least 1 transposon insertion in at least two out of the three replicates of the “total pool” samples. Also, the normalized abundance of insertions within a gene had to be greater than 0.01 in the “total pool” when averaged across all three replicates (where 1 is the expected normalized abundance for a “neutral” gene). These cutoffs allowed us to assess the phenotype of ~83% of the genes (2747/3310) in the *A. baylyi* ADP1 genome. For visualization, the relative abundance of Tn insertions within each gene is plotted from the “transformant pool” relative to the “total pool”.

### Pilin labeling, imaging, and quantification

Pilin labeling was performed as described previously with some changes^2,3^. Briefly, 100 μl of overnight-grown cultures was added to 900 μl of LB in 1.5 ml microcentrifuge tube, and cells were grown at 30 °C rotating on a roller drum for 70 min. Cells were then centrifuged at 18,000 x *g* for 1 min and then resuspended in 50 μl of LB before labeling with 25 μg/ml of AlexaFluor488 C5-maleimide (AF488-mal) (ThermoFisher) for 15 min at room temperature. Labeled cells were centrifuged, washed once with 100 μl of LB without disrupting the pellet, and resuspended in 5-20 μl LB. Cell bodies were imaged using phase contrast microscopy while labeled pili were imaged using fluorescence microscopy on a Nikon Ti-2 microscope using a Plan Apo 100X objective, a GFP filter cube for pili, a Hamamatsu ORCAFlash4.0 camera, and Nikon NIS Elements Imaging Software. Cell numbers and the percent of cells making pili were quantified manually using ImageJ^33^. All imaging was performed under 1% UltraPure agarose (Invitrogen) pads made with phosphate-buffered saline (PBS) solution.

### Western blotting

~10^9^ Cells from overnight cultures were concentrated into a pellet by centrifugation, and the culture supernatant was removed. Cell pellets were resuspended in 50 μl PBS and then mixed with an equal volume of 2×SDS-PAGE sample buffer (125 mM Tris, pH 6.8, 20% glycerol, 4% SDS, 0.4% bromophenol blue, and 10% β-mercaptoethanol) and boiled using a heat block set to 100 °C for 10-15 min. Proteins were separated on a 4-20% pre-cast polyacrylamide gel (Biorad) by SDS electrophoresis, electrophoretically transferred to a nitrocellulose membrane, and probed with mouse monoclonal α-FLAG antibodies (Sigma), mouse monoclonal α-GFP and/or mouse monoclonal α-RpoA (BioLegend) primary antibodies. Blots were then incubated with α-mouse IRDye secondary antibodies (Licor) and imaged using a Licor imaging system.

### Coimmunoprecipitation (pulldown) experiments

Overnight cultures of cells grown in tubes in LB medium were diluted by 1/10 into fresh LB for a total volume of 30 ml or 50 ml in 125 ml or 250 ml volume flasks respectively. 30-50 ml cultures were grown to exponential growth phase by shaking for 1.5 hr at 30 °C. The total culture volume was then harvested at 10,000 x *g* for 10 min at room temperature, and the supernatant was removed. Cell pellets were resuspended in 2 ml of Buffer 1 (50 mM Tris-Cl pH 7.4, 150 mM NaCl, 1 mM EDTA) and transferred to 2 ml volume microcentrifuge tubes and centrifuged at 18,000 x *g* for 1 min. Cells were washed once more with 2 ml of Buffer 1, and washed pellets were resuspended in 1 ml of Buffer 2 (50 mM Tris-Cl pH 7.4, 150 mM NaCl, 1 mM EDTA, 10 mM MgCl2, 0.1% Triton X-100, 2% glycerol). To lyse cells, 4200 units of Ready-Lyse lysozyme (Lucigen), 30 units of DNase I (New England Biolabs), and 10 μl of concentrated protease inhibitor cocktail (Sigma) (one pellet dissolved in 500 μl of Buffer 1) were added to cell suspensions and incubated at room temperature for 45 min. Cell debris was removed by centrifugation at 10,000 x *g* for 5 min at 4 °C. 50 μl of cell lysates were set aside and used as an “input” sample. 50 μl aliquots of α-FLAG magnetic bead slurry (Sigma) or α-GFP magnetic bead slurry (MBL biotech) in 1.5 ml microcentrifuge tubes were washed three times with 1 ml of Buffer 2 using a magnetic collection stand. 1 ml of cell lysates was added to washed magnetic beads and subjected to end-over-end rotation at 4 °C for 2 hr. Beads were then washed three times with 0.5 ml of Buffer 2, with 10 min incubations in Buffer 2 at 4 °C between each wash step. Beads were briefly washed a 4^th^ time with 0.5 ml Buffer 2. To elute proteins from α-FLAG beads, 100 μl of elution buffer (150 μg/ml 3X-FLAG peptide, Sigma, in Buffer 2) was added and samples were subjected to end-over-end rotation at 4 °C for 30 min. To elute proteins from α-GFP beads, beads were resuspended in 50 μl PBS, 2x SDS-PAGE sample buffer was added to tubes, and samples were boiled as described above. Eluates and input samples were subjected to western blotting as described above.

### Measuring GFP fluorescence in reporter strains

*V. cholerae* reporter strains harboring P_*cpiA*_-GFP were grown to late log in LB medium with or without 0.2% arabinose as indicated. Cells were then washed once in instant ocean medium (7g/L; Aquarium Systems) and transferred to a 96-well plate. Fluorescence was then determined on a Biotek H1M plate reader with monochromater set to 500 nm for excitation and 540 nm for emission.

### Fluorescent DNA binding/uptake

A ~7 kb PCR product was fluorescently labeled as described previously^11^ using the Cy3 LabelIT kit (Mirus Biosciences) as per manufacturer recommendations. For the parent strain, 1 μl (100 ng) of Cy3-DNA was added to 900 μl of LB in a 1.5 ml microcentrifuge tube along with 100 μl of overnight-grown cultures, and cells were grown at 30°C rotating on a roller drum for 70 min. Pili were then labeled as described above.

To visualize DNA-binding in the *pilT* mutant, cells were grown as described above, but 1 μl (100 ng) of Cy3-DNA was added to cells along with AF488-mal and incubated for 25 min before washing. Cells were imaged using the same microscopy setup described above, using a dsRed filter cube to image Cy3-DNA.

### Phylogenetic analysis

TfpB homologues were identified by BLAST using 43 manually selected Gammaproteobacteria genomes^34^ through Integrated Microbial Genomes and Microbiomes online resources (IMG)^35^ resulting in 209 protein sequences. The FtsK protein from *A. baylyi* ADP1 was added manually and used as an outgroup. Through the NGPhylogeny.fr server^36^, sequences were aligned using the default parameters of MAFFT^37^, and phylogenetic tree construction was performed on the aligned sequences using the default parameters of FastTree software^38,39^ set to perform 100 bootstraps^40^. The resulting tree was visualized using the Interactive Tree of Life (iTOL) visualization software^41^, from which trees were exported for publication.

## Acknowledgements

We would like to thank E. Geisinger for *A. baylyi* ADP1 ATCC33305 wildtype strain, and we would like to thank K. Hummels and B. Bratton for helpful suggestions on phylogenetic analysis. CKE is a Damon Runyon Fellow supported by the Damon Runyon Cancer Research Foundation (DRG-2385-20). This work was supported in part by the National Science Foundation, through the Center for the Physics of Biological Function (PHY-1734030). This work was supported by grant R35GM128674 from the National Institutes of Health awarded to ABD and the National Institutes of Health Pioneer Award 1DP1AI124669-01 awarded to ZG.

## Author contributions

CKE and ABD designed and coordinated the overall study. CKE, TND, CAK and ABD performed the experiments. All authors analyzed and interpreted data. CKE and ABD wrote the manuscript with help from ZG and JWS.

## Data availability statement

The data that support the findings of this study are available from the corresponding authors upon request.

## Supplemental Materials for

**Supplemental figure 1.**
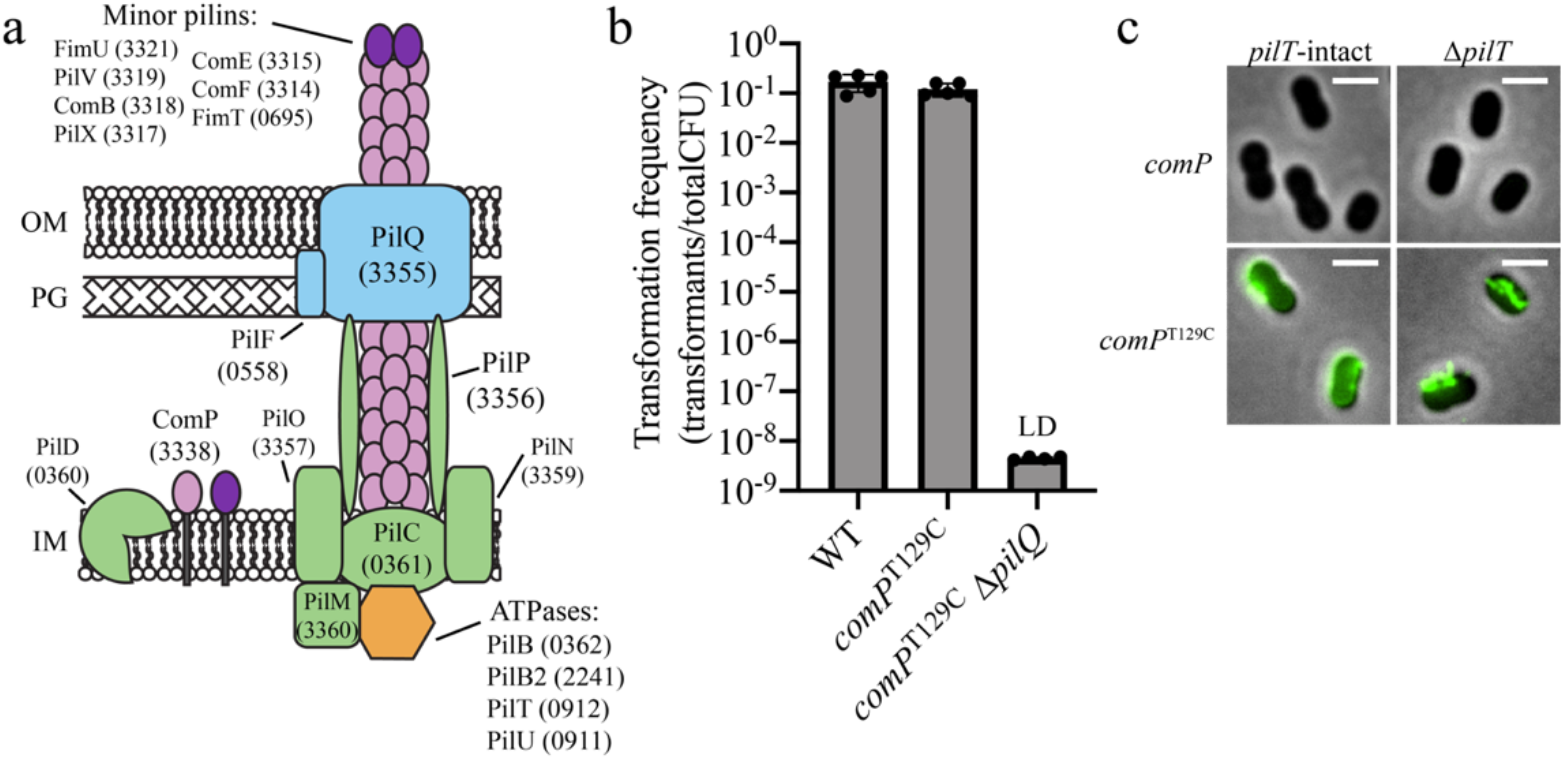
The strain for labeling T4P in *A. baylyi* is fully functional. (a) Schematic of assembled T4P components found in *A. baylyi*. Numbers in parentheses are ACIAD numbers associated with each component. OM, outer membrane; PG, peptidoglycan; IM, inner membrane. (b) Natural transformation assays of indicated strains. Each data point represents a biological replicate and bar graphs indicate the mean ± SD. The transformation frequency of the Δ*pilQ* strain was below the limit of detection, indicated by LD. (c) Representative images of indicated strains labeled with AF488-mal with background fluorescence subtracted. Scale bars, 2 μm.

**Supplemental figure 2.**
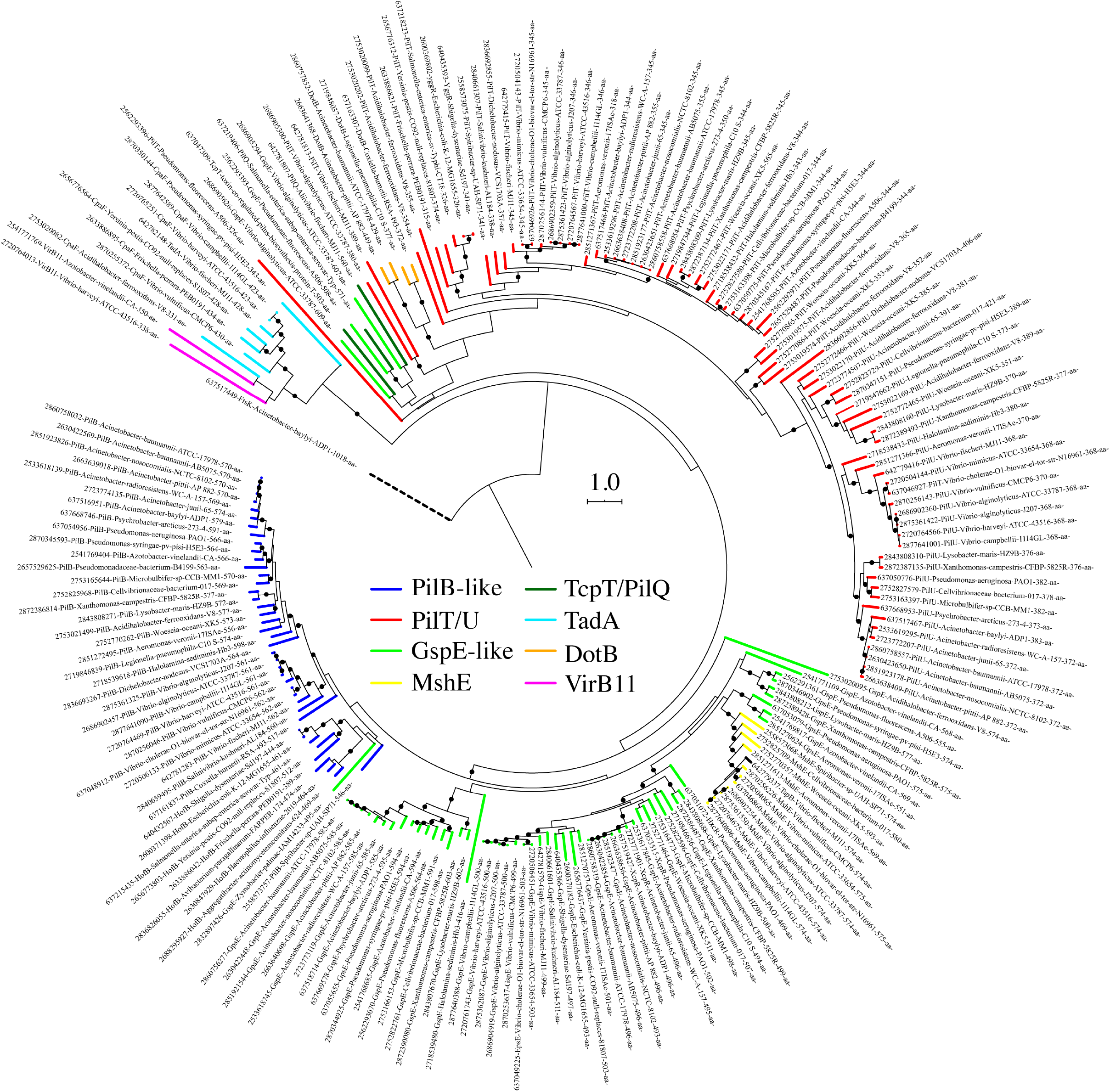
TfpB clusters with a group of other GspE-like proteins that are distinct from PilB proteins. A rooted phylogeny of TfpB homologues shown in Figure 2d with labels. Labels indicate IMG GeneID, annotated protein name, species, protein size. Nodes with bootstrap values greater than or equal to 70% are indicated by black circles.

**Supplemental figure 3.**
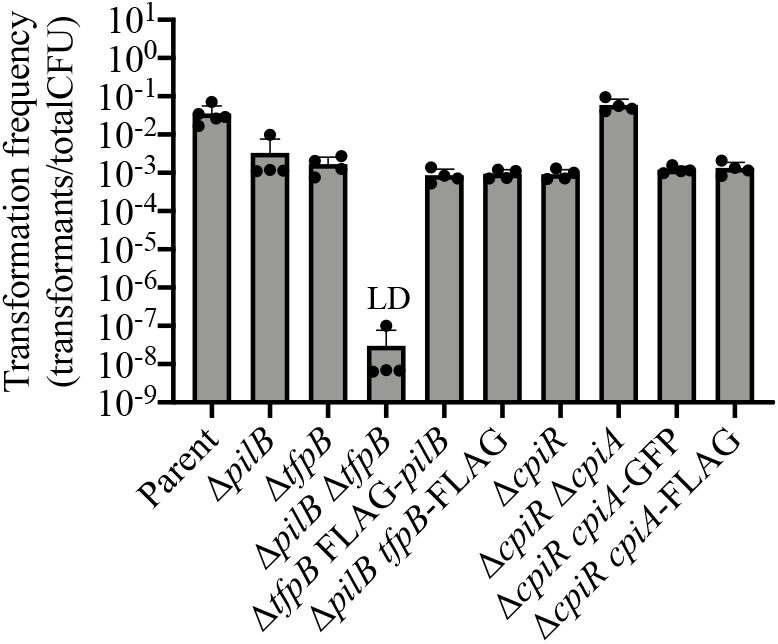
Natural transformation assays of indicated strains. Each data point represents a biological replicate and bar graphs indicate the mean ± SD. The transformation frequency of the Δ*pilB ΔtfpB* strain was below the limit of detection, indicated by LD.

**Supplemental figure 4.**
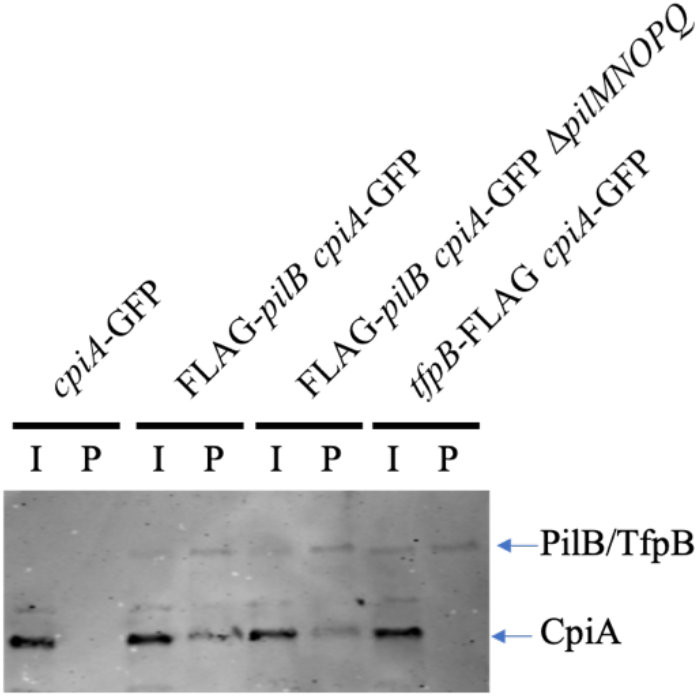
CpiA specifically interacts with PilB to inhibit its activity. Western blot showing coimmunoprecipitation experiments where FLAG-PilB or TfpB-FLAG were used as the bait proteins to test for interaction with CpiA-GFP as the prey protein in Δ*cpiR* mutant backgrounds. I, input sample; P, pulldown (coimmunoprecipitation) sample.

**Supplemental table 1.** Bacterial strains used in this study.

| CE# | ZG# | TND/<br>SAD# | Strain name in<br>manuscript | Genotype | Figure ('S'<br>denotes<br>supplemental<br>figure) |
| --- | --- | --- | --- | --- | --- |
| CE1 |  | SAD631 | Wildtype (WT) | *Acinetobacter baylyi* ADP1 | S1B, S1C |
| CE100 |  | SAD2537 | Pil-cys or<br>Parent | ADP1 *comP*<sup>T129C</sup> | 1A, 1C, 1D, 2A,<br>3B, S1B, S1C,<br>S3 |
| CE329 | ZG1709 | | $\Delta pilT$ | ADP1 $\Delta pilT$ | S1C |
| CE317 | | TND0779 | Pil-cys $\Delta pilT$ or<br>Parent | ADP1 *comP*<sup>T129C</sup> $\Delta pilT::spec$ | 1A, 1C, 2B, S1C |
| | | TND0114 | $\Delta comP$ | ADP1 $\Delta comP::kan$ | 2A |
| CE150 | ZG1680 | | $\Delta pilC$ | ADP1 *comP*<sup>T129C</sup> $\Delta pilC$ | 1C, 1D |
| CE16 | ZG1678 | | $\Delta pilY1$ | ADP1 *comP*<sup>T129C</sup> $\Delta pilY1$ | 1C, 1D |
| CE66 | ZG1679 | | $\Delta pilQ$ | ADP1 *comP*<sup>T129C</sup> $\Delta pilQ$ | 1C, 1D, S1B |
| CE286 | | TND2544 | $\Delta comEA$ | ADP1 *comP*<sup>T129C</sup><br>$\Delta comEA::kan$ | 1C, 1D |
| CE287 | | TND2546 | $\Delta comEC$ | ADP1 *comP*<sup>T129C</sup><br>$\Delta comEC::kan$ | 1C, 1D |
| CE289 | | TND2556 | $\Delta comM$ | ADP1 *comP*<sup>T129C</sup> $\Delta comM::kan$ | 1C, 1D |
| CE22 | ZG1683 | | $\Delta pilB$ | ADP1 *comP*<sup>T129C</sup> $\Delta pilB$ | 2A, 2C, 4C, S3 |
| CE67 | ZG1685 | | $\Delta tfpB$ | ADP1 *comP*<sup>T129C</sup> $\Delta tfpB$ | 2A, 2C, 4C, S3 |
| CE54 | ZG1684 | | $\Delta pilB \Delta tfpB$ | ADP1 *comP*<sup>T129C</sup> $\Delta pilB$<br>$\Delta tfpB::kan$ | 2A, 2C, 4C, S3 |
| CE310 | ZG1673 | | $\Delta pilB \Delta pilT$ | ADP1 *comP*<sup>T129C</sup> $\Delta pilB$<br>$\Delta pilT::spec$ | 2B, 2C, 4B |
| CE311 | ZG1674 | | $\Delta tfpB \Delta pilT$ | ADP1 *comP*<sup>T129C</sup> $\Delta tfpB$<br>$\Delta pilT::spec$ | 2B, 2C, 4B |
| CE316 | ZG1677 | | $\Delta pilB \Delta tfpB$<br>$\Delta pilT$ | ADP1 *comP*<sup>T129C</sup> $\Delta pilB$<br>$\Delta tfpB::kan \Delta pilT::spec$ | 2B, 2C |
| CE263 | | TND1201 | $\Delta cpiR$ | ADP1 *comP*<sup>T129C</sup> $\Delta cpiR::kan$ | 3B, 4A, S3 |
| CE264 | | SAD2792 | $\Delta cpiR \Delta cpiA$ | ADP1 *comP*<sup>T129C</sup> $\Delta cpiRA::kan$ | 3B, 4A, S3 |
|  |  | TND2260 | *P*<sub>tac</sub>-*cpiA* | ADP1 *comP*<sup>T129C</sup><br>igACIAD0096-ACIAD0098::*Cm*<sup>R</sup>-*P*<sub>tac</sub>-*cpiA* | 3B |
| | | TND2093 | *P*<sub>cpiA</sub>-GFP<br>Parent | *V. cholerae* E7946 *Sm*<sup>R</sup><br>$\Delta lacZ::SpecR$-*P*<sub>cpiA</sub>-*gfp* | 3C |
| | | TND2126 | *P*<sub>cpiA</sub>-GFP *P*<sub>BAD</sub>-<br>*cpiR* | *V. cholerae* E7946 *Sm* <sup>R</sup><br>$\Delta lacZ::SpecR$ - *P* <sub>cpiA</sub> - *gfp*<br>$\Delta VCA0692::CarbR-PBAD-cpiR$ | 3C |
|  |  | TND1335 | *cpiA*-FLAG | ADP1 *comP*<sup>T129C</sup> *cpiA*-3X<br>FLAG | 3D |
| | | TND1284 | *cpiA*-FLAG<br>$\Delta cpiR$ | ADP1 *comP*<sup>T129C</sup> *cpiA*-3X<br>FLAG $\Delta cpiR::kan$ | 3D, S3 |
| CE282 | | SAD2808 | $\Delta pilB \Delta cpiR$ | ADP1 *comP*<sup>T129C</sup> $\Delta pilB$<br>$\Delta cpiR::kan$ | 4A, 4C |
| CE283 | | SAD2806 | $\Delta tfpB \Delta cpiR$ | ADP1 *comP*<sup>T129C</sup> $\Delta tfpB$<br>$\Delta cpiR::kan$ | 4A, 4C |
| CE284 | | SAD2809 | $\Delta pilB \Delta cpiR$<br>$\Delta cpiA$ | ADP1 *comP*<sup>T129C</sup> $\Delta pilB$<br>$\Delta cpiRA::kan$ | 4A |
| CE285 | | SAD2807 | $\Delta tfpB \Delta cpiR$<br>$\Delta cpiA$ | ADP1 *comP*<sup>T129C</sup> $\Delta tfpB$<br>$\Delta cpiRA::kan$ | 4A, 4C |
| CE265 | | SAD2645 | $\Delta cpiR \Delta pilT$ | ADP1 $comP^{T129C} \Delta cpiR::kan \Delta pilT::spec$ | 4A, 4B |
| CE266 | | SAD2810 | $\Delta cpiR \Delta cpiA \Delta pilT$ | ADP1 $comP^{T129C} \Delta cpiRA::kan \Delta pilT::spec$ | 4A |
| CE302 | | SAD2822 | $\Delta pilB \Delta cpiR \Delta pilT$ | ADP1 $comP^{T129C} \Delta pilB \Delta cpiR::kan \Delta pilT::spec$ | 4A, 4B |
| CE303 | | SAD2820 | $\Delta tfpB \Delta cpiR \Delta pilT$ | ADP1 $comP^{T129C} \Delta tfpB \Delta cpiR::kan \Delta pilT::spec$ | 4A, 4B |
| CE304 | | SAD2823 | $\Delta pilB \Delta cpiR \Delta cpiA \Delta pilT$ | ADP1 $comP^{T129C} \Delta pilB \Delta cpiRA::kan \Delta pilT::spec$ | 4A |
| CE305 | | SAD2821 | $\Delta tfpB \Delta cpiR \Delta cpiA \Delta pilT$ | ADP1 $comP^{T129C} \Delta tfpB \Delta cpiRA::kan \Delta pilT::spec$ | 4A, 4B |
| | | TND2904 | $\Delta tfpB$ FLAG-$pilB$ | ADP1 $comP^{T129C} \Delta tfpB::kan$ 3X FLAG-$pilB$ | S3 |
| | | TND2892 | $\Delta pilB$ $tfpB$-FLAG | ADP1 $comP^{T129C} \Delta pilB::kan$ $tfpB$-3X FLAG | S3 |
| | | TND1296 | $cpiA$-GFP $\Delta cpiR$ | ADP1 $comP^{T129C} cpiA$-GFP $\Delta cpiR::kan$ | S3, S4 |
| | | TND2337 | FLAG-$pilB$ | ADP1 $comP^{T129C}$ 3X FLAG-$pilB$ | 3D |
| CE608 | ZG1784 | | FLAG-$pilB \Delta cpiR$ | ADP1 $comP^{T129C}$ 3X FLAG-$pilB \Delta cpiR::kan$ | 3D, 4D |
| | | TND2383 | FLAG-$pilB cpiA$-GFP $\Delta cpiR$ | ADP1 $comP^{T129C}$ 3X FLAG-$pilB cpiA$-GFP $\Delta cpiR::kan$ | 4D, S4 |
| CE626 | ZG1785 | | FLAG-$pilB cpiA$-GFP $\Delta cpiR \Delta pilMNOPQ$ | ADP1 $comP^{T129C}$ 3X FLAG-$pilB cpiA$-GFP $\Delta cpiR::kan \Delta pilMNOPQ::zeo$ | S4 |
| CE656 | ZG1786 | | FLAG-$tfpB cpiA$-GFP $\Delta cpiR$ | ADP1 $comP^{T129C} tfpB$-3X FLAG $cpiA$-GFP $\Delta cpiR::kan$ | S4 |

CE strains available upon request from CKE, ZG strains available upon request from ZG, TND or SAD strains available upon request from ABD.

**Supplemental table 2.**
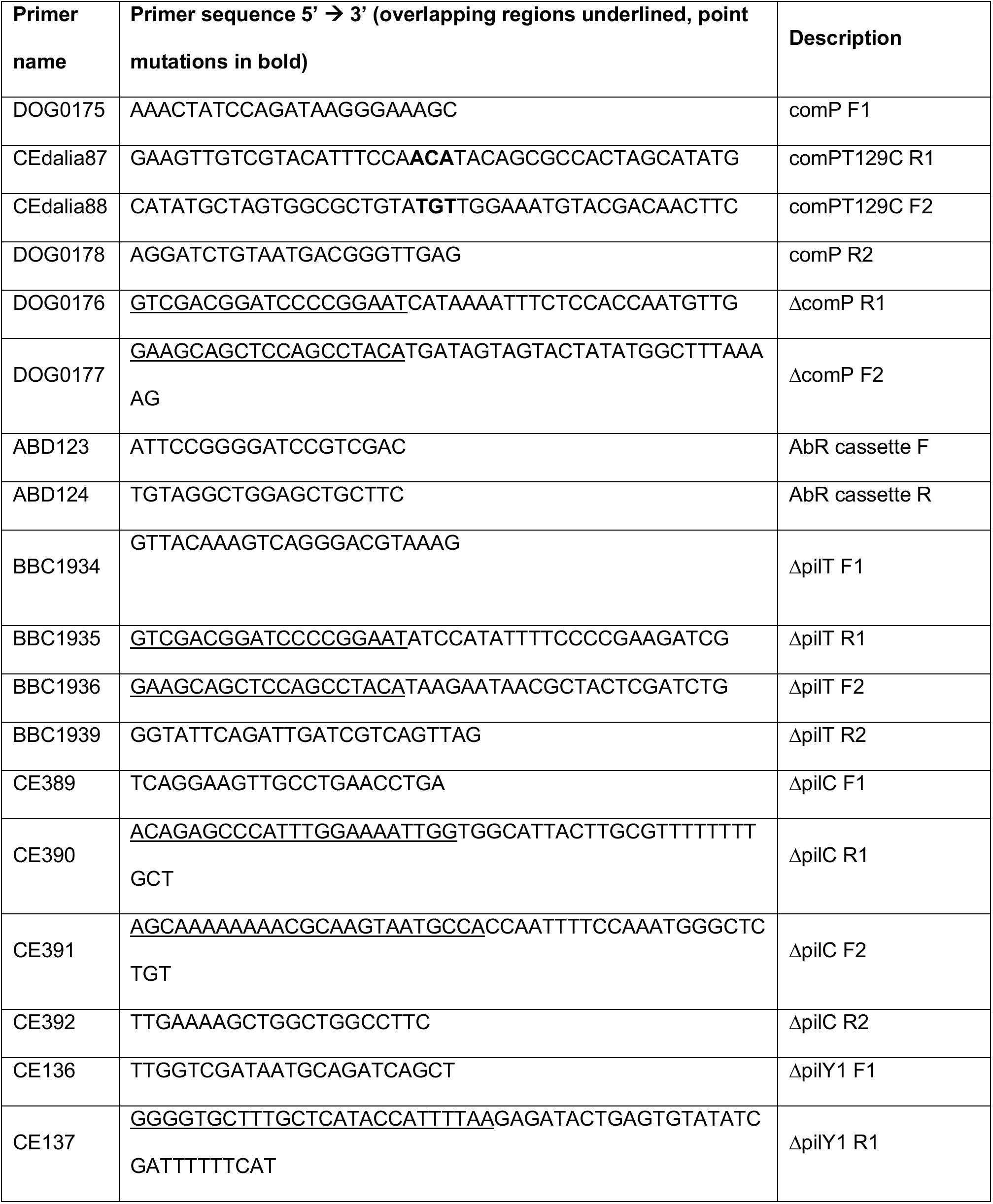
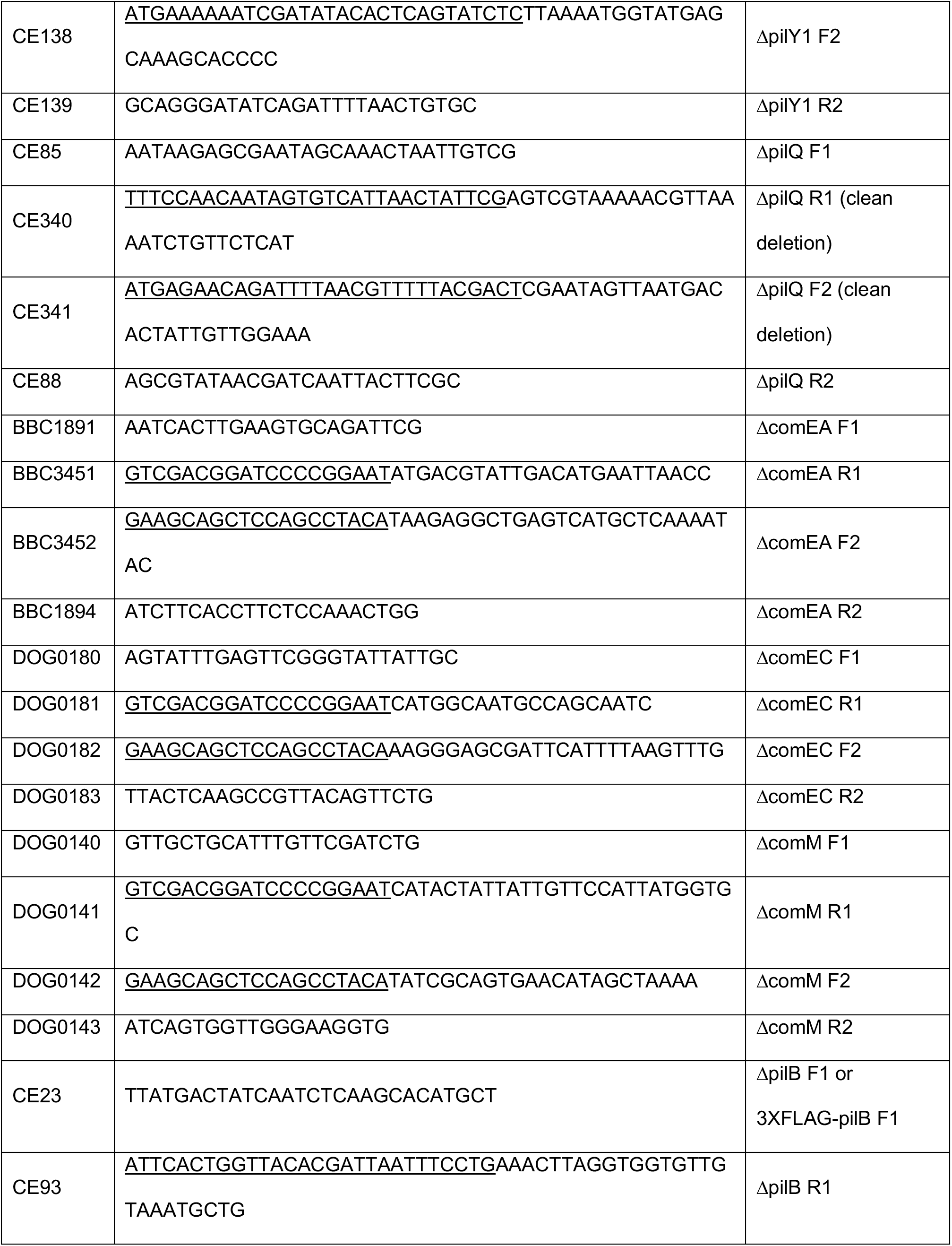
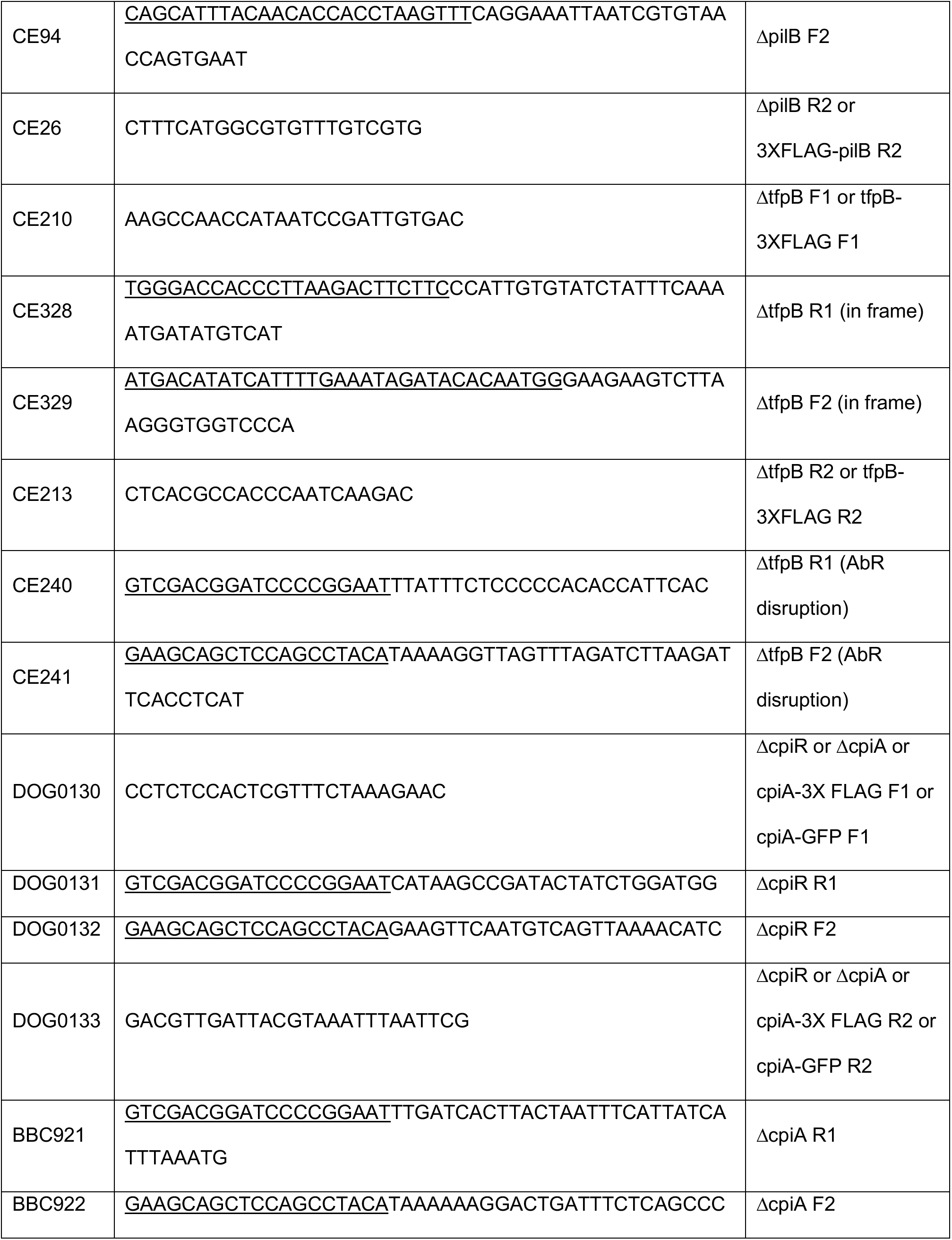
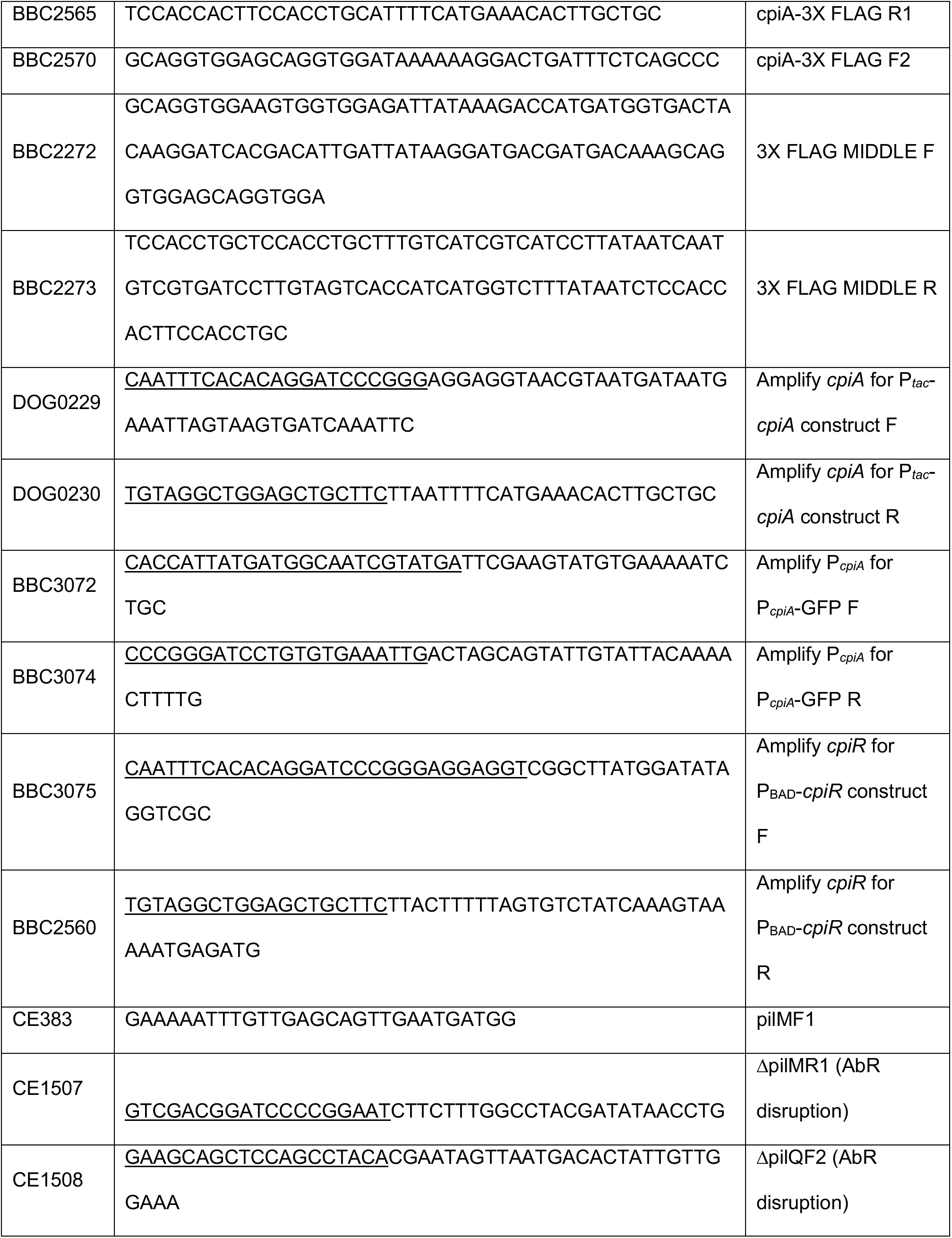
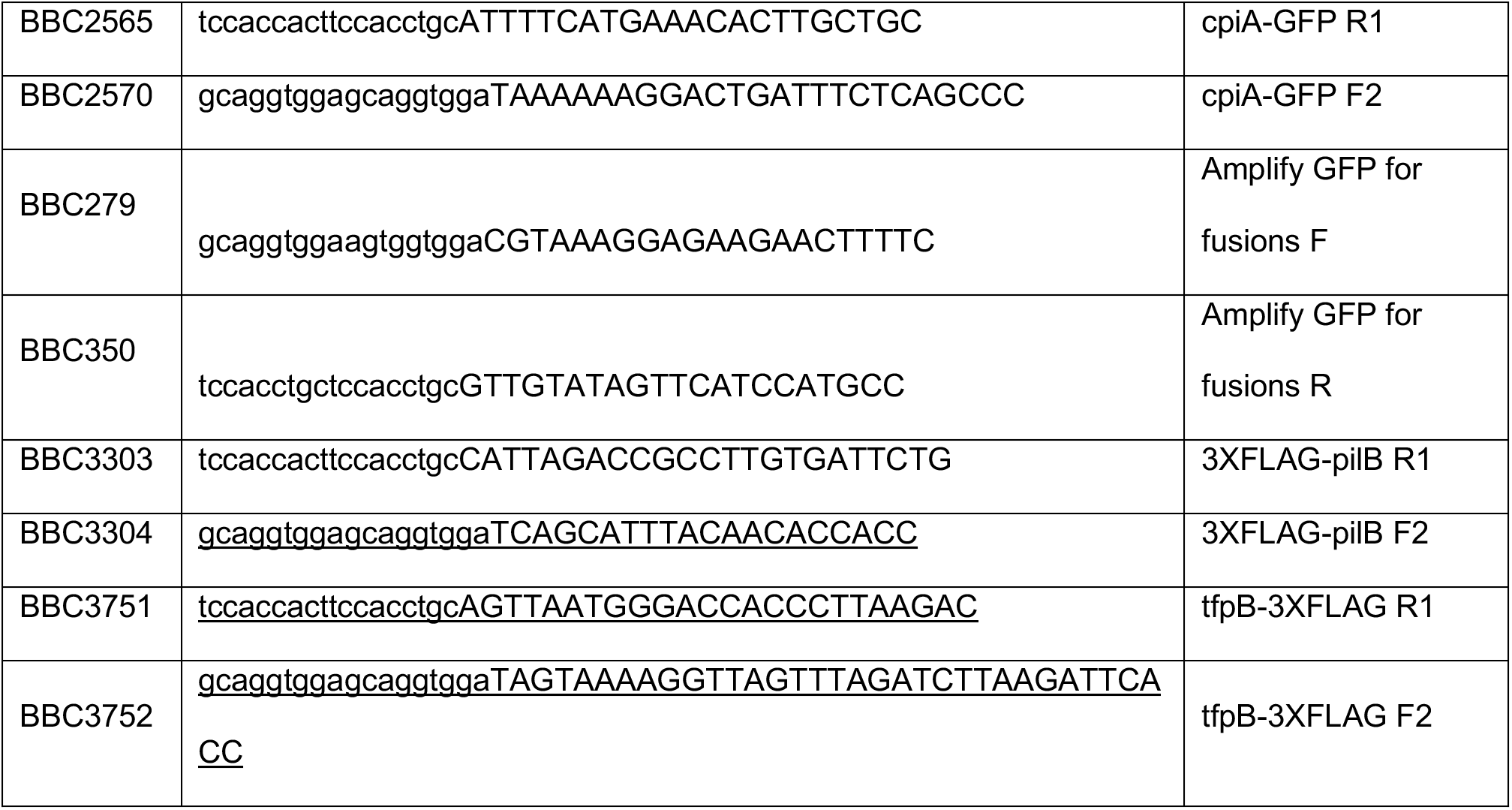
Primers used for strain construction.

